# Mapping of Shore Area Wetlands in Lake Tana Biosphere Reserve, Ethiopia Using Sentinel-1A SAR and Multi-Source Data

**DOI:** 10.1101/2024.12.30.630805

**Authors:** Yirga Kebede Wondim, Ayalew Wondie Melese, Workiyie Worie Assefa

## Abstract

Shore area wetlands (lacustrine fringe) play a critical role as ecotones that support biodiversity, provide habitats for spawning and refuge, and exhibit high levels of primary productivity. They facilitate significant exchanges of materials between aquatic and terrestrial ecosystems. To effectively manage and preserve these important resources, it is essential to understand their distribution, size, and dynamic changes. This study aimed to create an accurate map of shoreline wetlands using multi-temporal and multi-source data, wetland indicators (hydrology, vegetation, hydric soils), and radar imagery from Sentinel-1A, employing Geomatica software. Additionally, ArcGIS software was used to map the topographic position, Lake Bathymetry, and hydric soil indicators for wetlands. The analytical hierarchy process and weighted overlay methods were also applied in the mapping process for integrating all the indicators to obtain the final extent of shoreline wetlands. The topography position wetland indicator map covered about 55,364ha, while hydric soils covered around 55,151 ha within a 3 km buffer from Lake Tana. The map of hydrology indicator for wetlands revealed that permanently inundated areas accounted for roughly 591,312 ha, and when temporarily inundated areas were included, the total coverage increased to 607,053 ha. Hydrophytic vegetation, including invasive water hyacinth, covered over 74,772 ha. Overall, shoreline wetlands were predominantly located within three kilometers of the terrestrial area from Lake Tana, totaling 26,664 ha. The overall accuracy of land use and cover classification was recorded at 79%, with a Kappa statistic of 0.70, indicating that the resulting map is of acceptable quality. The integration of multi-temporal and multi-source data, along with wetland indicators and radar imagery from Sentinel-1A using Geomatica software, has provided valuable insights into the spatial distribution of shoreline wetlands in Lake Tana. The findings from this study will serve as an important reference for future research aimed at effectively managing and conserving these vital resources.

## Introduction

Shore area wetlands (lacustrine fringe) are important ecotones for biodiversity conservation, providing spawning and refuge habitats, exhibiting high primary productivity, and facilitating significant material exchange between aquatic and terrestrial ecosystems [1]. Despite their importance, lacustrine wetlands are among the most vulnerable ecosystems to human activities, particularly due to water level fluctuations [2]. Lake Tana, the largest freshwater lake in Ethiopia, is rich in lake-associated shoreline wetlands. These wetlands support spawning and juvenile habitats for several important fish species, including the unique endemic *Labeobarbus* [3], as well as a refuge for zooplankton [4]. Moreover, they serve as the feeding and breeding ground for various bird species such as the *common crane*, *wattled crane*, and *crowned crane* [5]. However, the wetlands of Lake Tana have faced severe threats from multiple anthropogenic activities; including recession agriculture, water level fluctuations, and siltation. The invasion of the lake by *Eichhornia crassipes* (water hyacinth) since 2011 has further exacerbated these challenges, as it quickly invades and dominates native wetland plant species [6]. Thus, it is essential to understand the distribution, size, and dynamic changes of the shoreline wetlands to effectively manage and preserve these crucial resources.

The use of GIS and remote sensing technologies for mapping and monitoring wetland resources has been well-established [7,8]. Remote sensing technology has been utilized to map wetland areas or changes in land use/cover in wetlands during the past five decades [9–11]. However, wetlands pose unique challenges for mapping due to their dynamic nature, environmental factors, and the limitations of traditional techniques [12]. Wetlands can be classified into various categories based on their hydrology, soils, water chemistry, vegetation, landscape position, and interests of the classifier. Most of the remote sensing techniques cannot detect water level changes beneath the vegetation in wetland ecosystems. Recently, the Synthetic Aperture Radar (SAR) technology has been applied effectively in conditions of persistent cloud, smoke, and haze, enabling the detection of water level changes across different wetland environments [13,14]. As an active microwave sensor, SAR can operate day or night, capturing images regardless of weather conditions [15]. This technology allows for the detection of water level changes under various weather situations by leveraging the ’double bounce’ phenomenon [16,17].Despite these advantages, speckle noise in SAR images reduces the image’s radiometric quality and complicates processing [18,19]. Fortunately, [20] demonstrated that utilizing an effective despeckling method can positively impact the precision of wetland classification.

The use of SAR data for wetlands studies has increased tremendously. Various sensors have been used in wetland investigations, including the ALOS Phased Array L-band Synthetic Aperture Radar (PALSAR), European Remote Sensing (ERS-1), RadarSAT, ASAR, Japanese Earth Resources Satellite 1 (JERS-1), AIRSAR, and TerraSAR-X Sentinel-1[21]. Wetland hydrology in areas such as the Everglades [16], Louisiana, and the Sian Ka’an in Yucatan [22] has been effectively assessed using Interferometric SAR (InSAR) observations. The same technique has been applied for multi-temporal monitoring of wetland water levels in the Florida Everglades (16), and has been combined with Radarsat-1 data to map fluctuations in water levels in coastal wetlands in southeastern Louisiana [15]. However, the use of radar data for the study of land use/cover, particularly wetlands, remains quite limited in Ethiopia. Only one study has been conducted on mapping the Dabus wetlands in Ethiopia by using Random Forest Classification of Landsat, PALSAR, and Topographic Data [11]. Additionally, a combination of sentinel-1 SAR and sentinel-2 MSI data has been employed for accurate urban land use-land cover classification in Gondar City [23].

Nevertheless, to our knowledge, there are currently no studies exploring the application of Sentinel-1A SAR for mapping shore area wetlands. Hence, this work aims to develop an accurate mapping of lacustrine wetlands using multi-temporal and multi-source data, indicators for wetlands, and radar imagery from Sentinel-1, all processed using Geomatica software. Specifically, this study will integrate spatial geo-information derived from topographic position and hydric soil indicators (essential for wetland identification) using ArcGIS software with data captured by the active sensor of Sentinel-1A SAR, which includes information on hydrophytic vegetation and wetland hydrology. The specific research components of this study are as follows: (a) Mapping topographic position (including Digital Elevation Models and its derivatives, as well as lake bathymetry) and hydric soil wetland indicators using various multi-source datasets and ArcGIS software; (b) Mapping hydrophytic vegetation and wetland hydrology using Sentinel-1 SAR data by processing in the Geomatica Banff software package; (c) Finally, integrating all wetland indicators to map shore area wetlands in Lake Tana Biosphere Reserve.

## Materials and Methods

### Description of the study area

Lake Tana (Fig.1) is the largest freshwater lake in Ethiopia. It is characterized by extensive shorelines and terrestrial wetlands [24], which provide essential spawning habitats for approximately half of the *Labeobarbu*s species, as well as three other commercially important fish species: Nile tilapia (*Oreochromis niloticus*), African catfish (*Clarias gariepinus*), and Beso (*Varicorhinus beso*)[3]. The juveniles of these fish species feed and grow in the lakeshore as well during the early years of their life [3], while the wetlands also serve as a refuge for zooplankton [4]. Additionally, these areas act as breeding grounds for various bird species, including the common crane (a wintering species), the endangered wattled crane, and the crowned crane[5].The lake and its associated wetlands provide immense socioeconomic benefits, supporting the livelihoods of thousands of people and contributing to economic activities through hydropower production and irrigation [25]. Despite their importance, the wetlands, including Lake Tana, have faced challenges from both short- and long-term water level fluctuations and heavy sediment loads, threatening their sustainable use.

**Fig.1.**
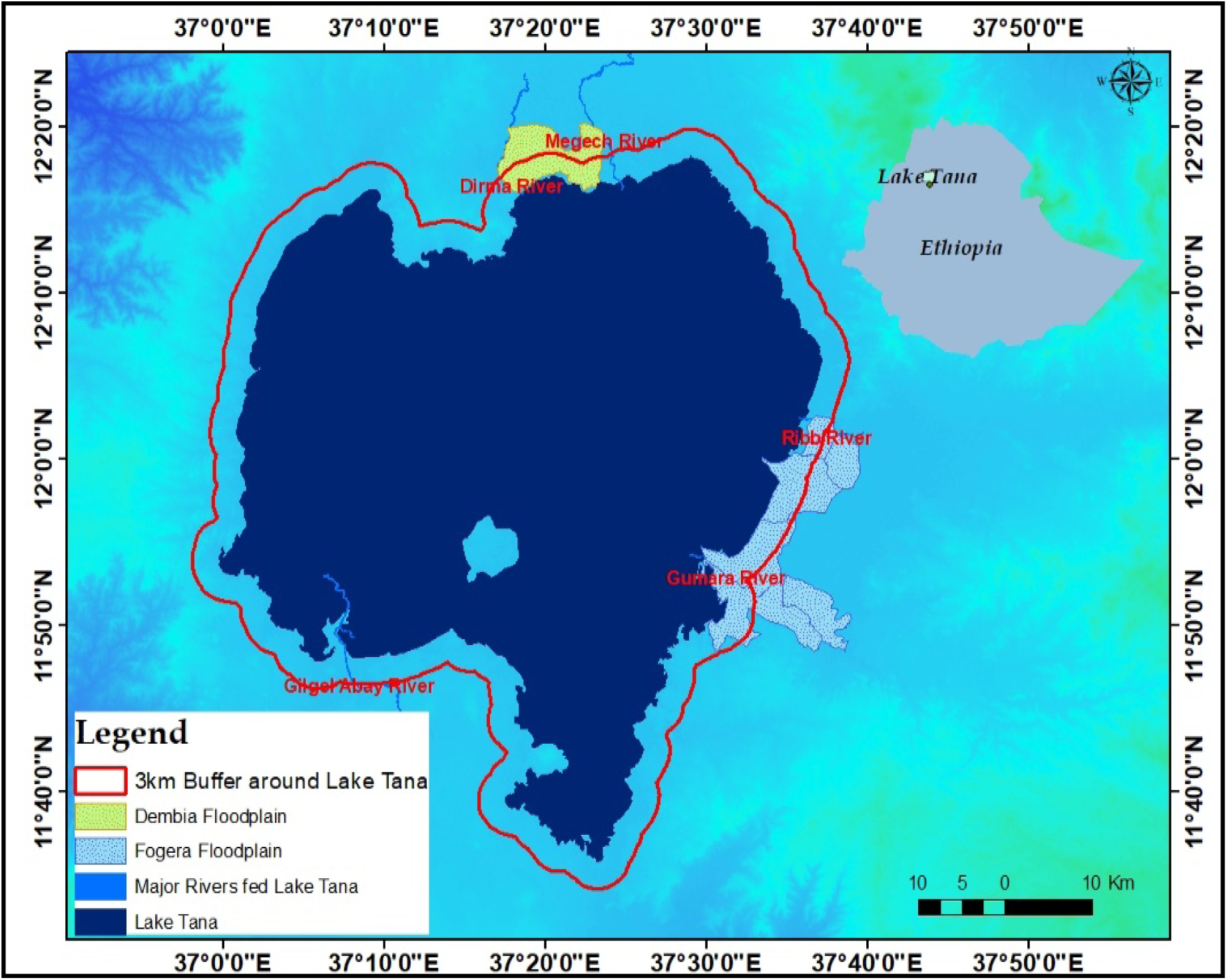
Location map of the study area

The shore area wetlands surround the lake (Fig. 1) and are formed and maintained through lacustrine processes. At mean annual water depths of less than 2 meters, these wetlands experience either permanent or periodic flooding. Tana, an adjacent freshwater lake, supplies water to these wetlands. In addition to the lake water, precipitation, and surface inflows also contribute to these lacustrine fringe wetlands. During the rainy season, the lake and rivers, along with precipitation, serve as the primary water inputs to the wetlands[24].

Within the shore area wetlands of Lake Tana, including the Lake itself, several physical, chemical, and biological processes work to purify the inflowing water and buffer the lake against pollution.[26] made the following generalizations after relating the circulation patterns of Lake Tana to the distribution of sediment, nutrients, and water hyacinth: (a) water, sediment, and associated nutrients, such as phosphorus generated from the eastern and northern catchments, had the longest retention time in the northeast of Lake Tana and as a result, most of the water is lost through evaporation while the sediment and phosphorus are partly suspended in the area and absorbed in the lakebed in the northeast, and (b) sediments and nutrients delivered from Gilgel Abay, southwest of Lake Tana, flow out of the lake through the two outlets in a shorter retention time favored by the close locations with the outlets.

### Mapping of shore area wetlands in Lake Tana

#### Mapping of wetland indicators

We utilized advanced technology by integrating multi-temporal and multi-source data for the classification, delineation, and mapping of the lacustrine wetlands at Lake Tana. This involved employing a range of automated, semi-automated, and manual techniques. For the automated approaches, key inputs included major soil types (Hydric Soils), Digital Elevation Models (DEM) and their derivatives, Lake Bathymetry, DEM/SRTM data, Synthetic Aperture Radar (SAR), and high-resolution Google Hybrid satellite imagery obtained through SAS Planet software. In addition to these automated methods, we conducted on-screen digitization of the enhanced imagery, manual delineation based on field data, and the delineation, merging, and naming of wetland parcels in accordance with political boundaries. Among the three primary wetland classification systems—Ramsar International Wetland classification, the Cowardin et al. system, and the hydrogeomorphic (HGM) classification [27,28] —we adapted the HGM system for this study. This approach focuses on three key factors: landscape position, major water supply, and hydrodynamics [28], which were essential for mapping and monitoring changes in the lacustrine fringe wetlands of Lake Tana. The general workflow of mapping shore area wetlands presented in the Fig.2.

**Fig.2.**
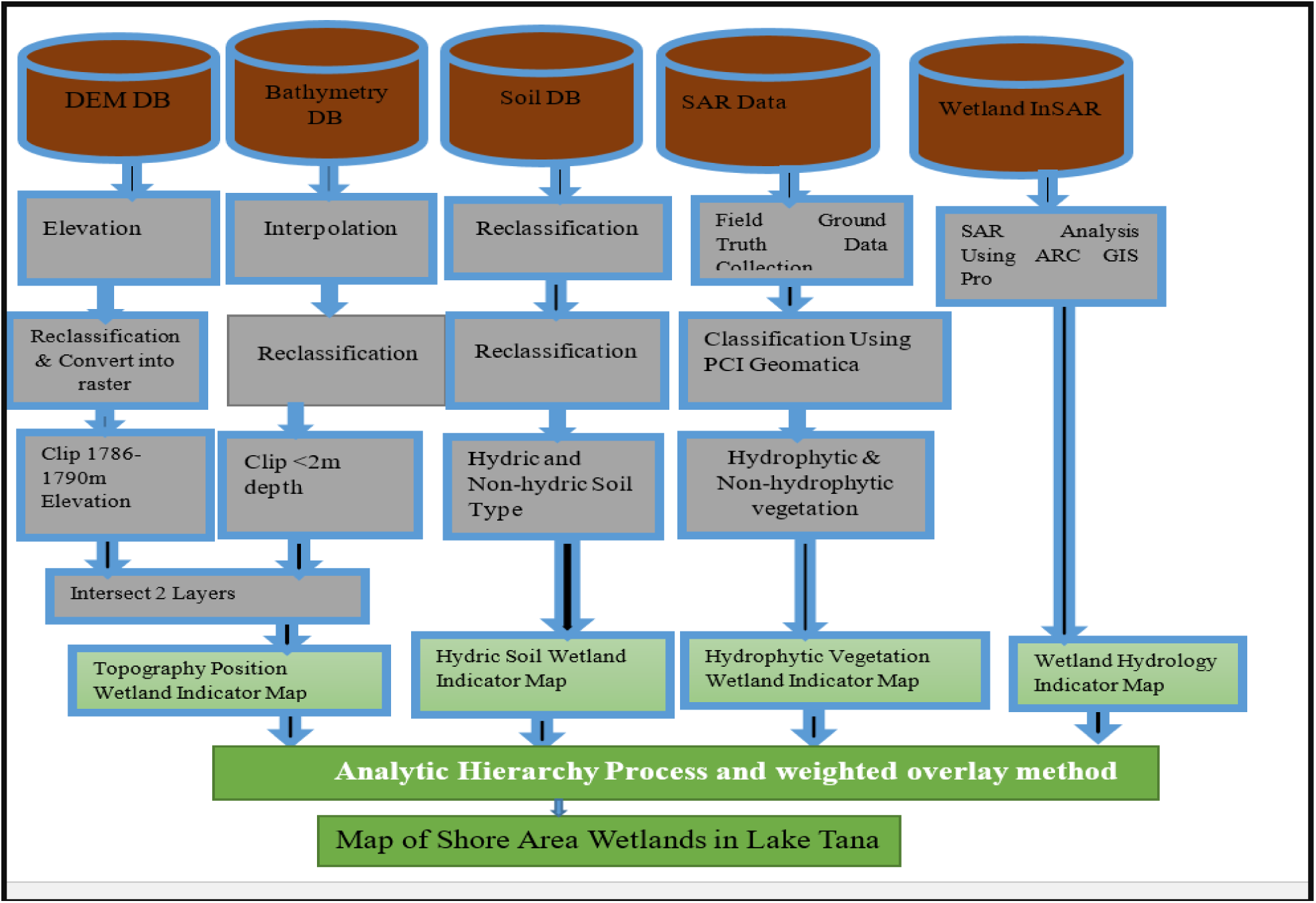
General workflow of mapping shore area wetlands

#### Topographic Position

Lacustrine fringe wetlands within the Lake Tana landscapes are created and sustained by lacustrine processes, primarily receiving water from the lake itself. Consequently, topographic position, particularly elevation, plays a crucial role in the formation of these wetlands. In this study, topographic position (DEM and its derivatives), and lake bathymetry are considered as an important indicator for mapping wetland areas along the shore. Bathymetric maps of wetlands serve many purposes, including defining legal boundaries, estimating water storage capacity and hydroperiod (depth and timing of flooding), as well as aiding in wetland design, restoration, and land use planning [29].

Secondary data from the bathymetry of Lake Tana during 2017 [30] was utilized to delineate the boundaries of the wetlands from the lake-ward side. In contrast, the average maximum lake level and elevation derivatives from the Digital Elevation Model (DEM) were employed to identify the boundaries of these wetlands from the landward side. Elevation values critical to Lake Tana and its surrounding areas, extending up to 1790 meters within a 3-kilometer buffer, were selected as they were expected to fall within the lacustrine wetland extending landward. The choice of 1790 meters as the critical elevation was based on the maximum lake level and additional water sources from nearby rivers.

The process of generating Topographic Position Mapping involved several key steps (Fig.3): (a) Reclassifying elevation maps into five categories within a 3 km buffer zone around the lake; (b) Identifying critical elevation values for Lake Tana and its surroundings, specifically up to 1790 meters within the 3 km buffer, to delineate the boundaries of shoreline wetlands from the landward side; (c) Reclassifying the lake bathymetry map into eight categories; (d) Utilizing the first category (0-2 meters depth) from the bathymetry map to define the boundaries of shoreline wetlands from the lake-ward side; and (e) Merging two layers—the elevation map for the range of 1786– 1790 meters and the bathymetry map for depths of 0-2 meters—to create the Topography Position Wetland Indicator Map.

**Fig. 3.**
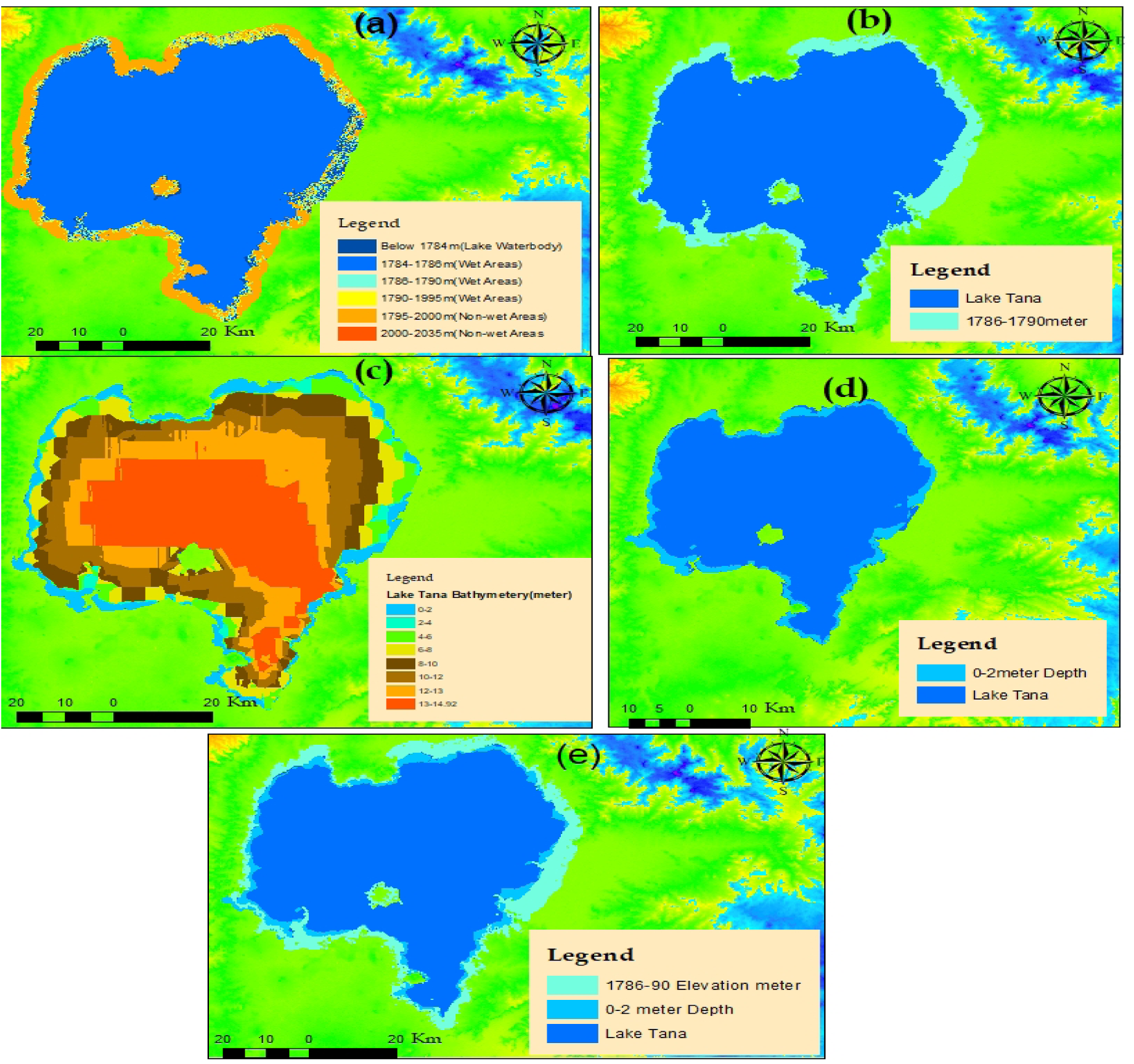
Topographic position mapping processes: (a) *elevation maps reclassified into 5 classes around 3km buffer zones of Lake Tana; (b) the critical elevation values of Lake Tana and the surrounding areas up to 1790 meters within the 3km buffer of Lake Tana, which were used to extract the boundaries of the lacustrine wetlands from the landward side; (c) Lake Tana bathymetry map reclassified into 8 classes; (d) from the 8 classes of bathymetry map, the first class (0–2 meter depth) was used to extract the boundaries of the lacustrine wetlands from the lake-ward side; and (e) two layers (c and d) were merged to produce the Topography Position Wetland Indicator Map*.

#### Hydric Soils

Hydric soils are characterized by saturation, ponding, or flooding for a duration sufficient during the growing season to foster the development of anaerobic conditions in the upper layers (31). These conditions create an ideal environment for the growth and reproduction of hydrophytic vegetation. In contrast, non-hydric soils are generally less conducive to the formation of wetlands. However, it is important to note that the absence of hydric soil does not necessarily indicate that wetlands are absent from the area [31]. Thus, to identify areas with a higher likelihood of wetland presence in this study, the hydric soil indicator was utilized alongside indicators of hydrophytic vegetation and topographic position.

Thus, to delineate hydric soils accurately, it was essential to employ national or regional field indicators of hydric soils or conduct a comprehensive soil survey. However, conducting detailed soil surveys for wetland mapping in this research project proved prohibitively expensive. As a result, we adopted a reference soil group map of Ethiopia [32], based on legacy data and machine learning techniques (EthioSoilGrids 1.0), which is the first of its kind for Ethiopia, along with a digital format of the Tana sub-basin soil survey for this study[33].

Three commonly recognized morphological traits can be utilized to differentiate between hydrophilic and non-hydrophilic soils: the presence of organic matter, gleying, and mottling or redoximorphic structures (31). By considering these established morphological characteristics, the major soil types surrounding Lake Tana within a 3 km buffer (Fig.4a) have been reclassified into a principal category labeled hydric, which includes Vertisols, Fluvisols, Gleysols, and Water. The remaining areas, comprising urban built-up zones, have been classified as non-hydric soils. Furthermore, these hydric soils are subdivided into water bodies, permanently hydric soils (Gleysols), and seasonally hydric soils (Vertisols and Fluvisols) (Fig.4b).

**Fig.4.**
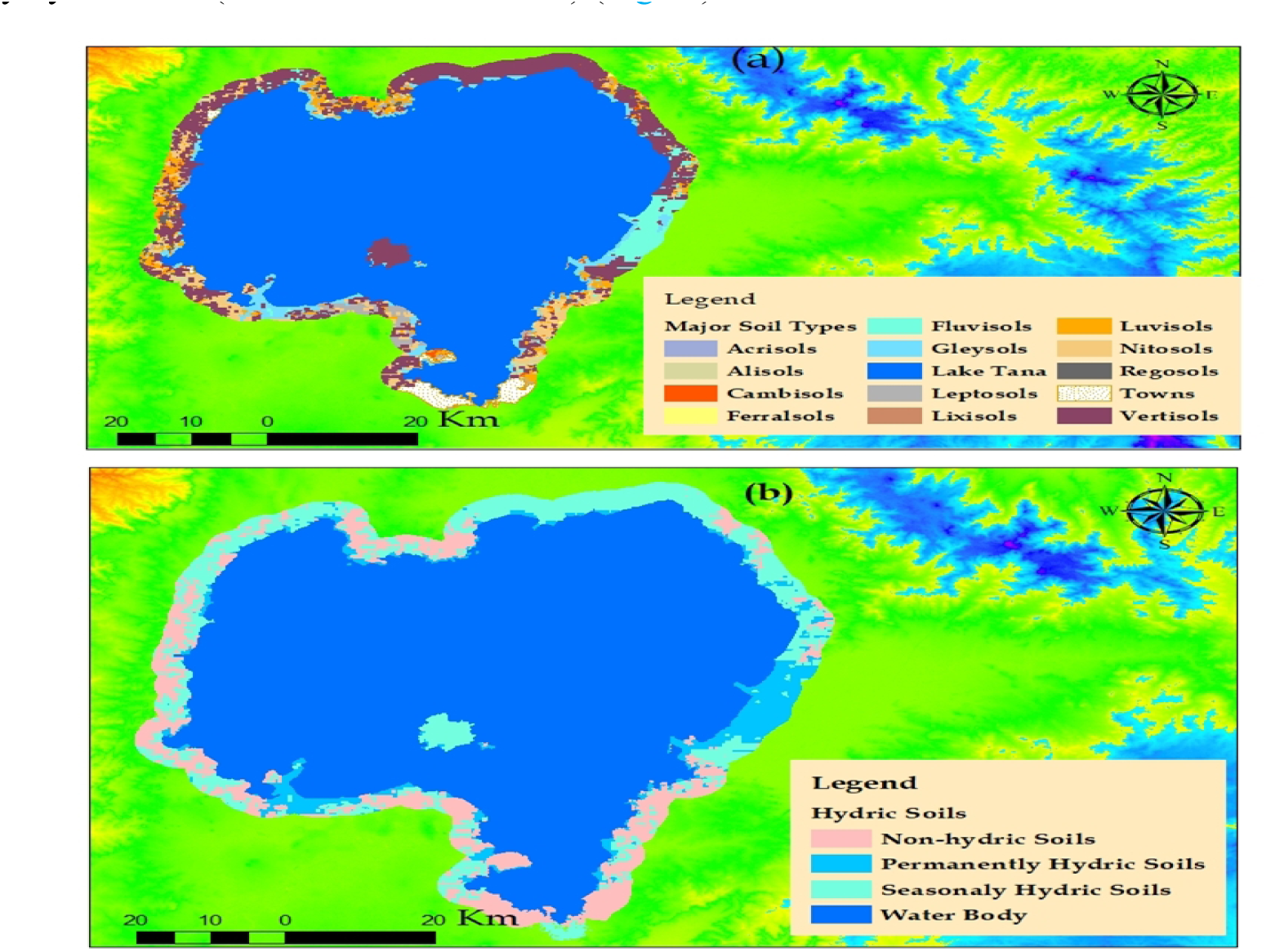
Hydric soil mapping processes: *Major soil types around Lake Tana up to a 3 km buffer extracted from a soil survey done by ADSWE (2014) (4a); and major soil types reclassified into water bodies, water bodies, permanently hydric soils (Gleysols), and seasonally hydric soils (Vertisols and Fluvisols) (4b)*.

#### Mapping Hydrophytic Vegetation Using SAR Data

Wetlands are characterized by their distinctive hydrophytic vegetation [34]. To enhance the efforts of wetland delineation and mapping in the study area, a survey of hydrophytic vegetation was conducted and integrated into the mapping process. By utilizing data collected across multiple seasons, we can improve the differentiation of wetland types through an analysis of seasonal variations in phenology that reflect changes in plant structure. This study specifically utilized wetland vegetation data gathered in May 2021 and October 2021.

In this study, hydrophytic vegetation was mapped as a wetland indicator using Sentinel-1A SAR data in conjunction with Object Analyst, an add-on package for PCI Geomatica Banff software. Sentinel-1A SAR was chosen for mapping hydrophytic vegetation in the shore area wetlands of Lake Tana for several reasons: (1) Sentinel-1A SAR is particularly effective when the performance of optical sensors is compromised by cloud cover and varying day/night conditions [15]; (2) The SAR signal’s ability to penetrate through vegetation and soil provides additional information that is not accessible through optical remote sensing data [12,19].

Within Geomatica Focus, Object Analyst provides a comprehensive interface that encompasses processes for segmenting imagery, extracting features, building training sites, classifying data (including the creation of classification rules), reshaping, and assessing accuracy (refer to Fig. 5a & 5b). Among the preprocessing steps, mosaicking was performed using the Mosaic Tool, while clipping was executed through Focus, Tools: Clipping/Sub-setting. The image was segmented into statistically homogeneous areas or objects (segments) that are uniform within themselves and distinct from nearby segments, utilizing image segmentation techniques (Fig.5c). Following segmentation, various attributes were calculated, including statistical, geometrical, and textural metrics. A ground truth image was subsequently created, with the collected training samples serving purposes for both supervised classification and accuracy assessment in Object-Based Image Analysis (OBIA). Ground truth points were gathered through field visits and SAS Planet imagery, resulting in ground truth shape-files overlaid with Sentinel-1A SAR images. Using these inputs, training site polygons were developed for both training and accuracy assessment (illustrated in Fig.5d). The classification function categorizes the segmented images into different land cover types based on the training dataset. Supervised classification was carried out using the Support Vector Machine technique [35,36].The advantage of this technique lies in its requirement for a minimal number of training samples [37]. Support Vector Machines (SVMs) are particularly effective for nonlinear classification challenges, which proves beneficial when extracting feature vectors from completely polarimetric SAR data. In this context, the radial basis function kernel was employed for classification, utilizing a high probability threshold of 0.9 and a penalty parameter of 100 [35]. The classification quality was assessed based on various metrics, including user’s accuracy, producer’s accuracy, overall accuracy (OAA), and the Kappa index of agreement (KIA) [38]. Following the classification process, the land cover classification results were updated, and an accuracy assessment was conducted [38]. The Fig.5 illustrates the workflow of the object-based image analysis (OBIA) process.

**Fig.5.**
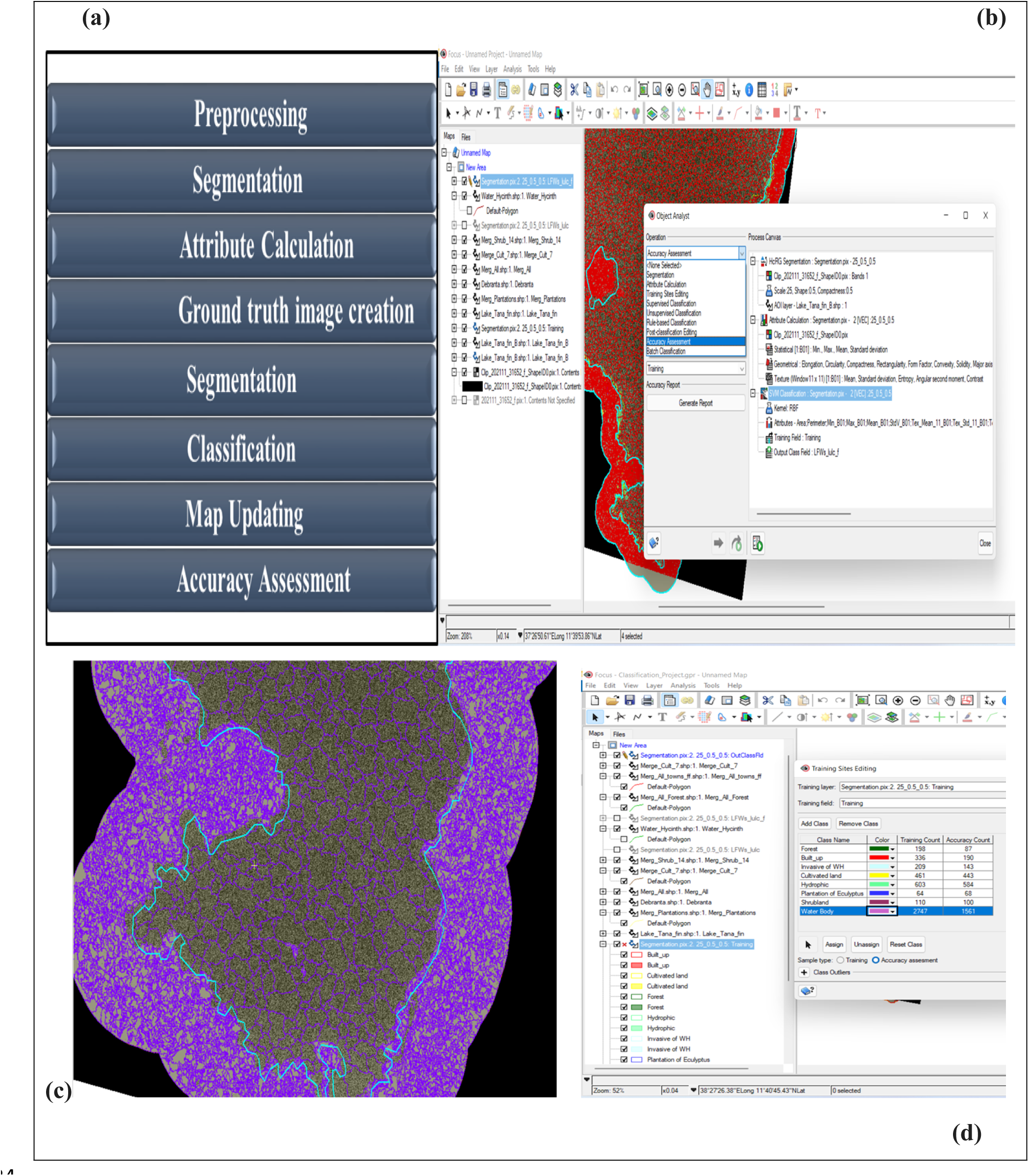
Process flow for object-based image analysis (OBIA): General workflow of hydrophytic vegetation mapping using object analyst within Geomatica Focus (5a), Process Canvas of Object Analyst (5b), Segmentation Result of Southern part Lake Tana including 3 km Buffer (5c) and Training Site Editing including Training Count and Accuracy Count (5d).

#### Wetland Hydrology Mapping Using SAR Data

Water determines the existence, establishment, and maintenance of specific types of wetlands and their associated activities (34).The distinct backscattering mechanisms exhibited by flooded plants and water enable the delineation of water bodies from surrounding vegetation [39].The varying roughness between water-covered areas and other land-cover types serves as the foundation for detecting surface waters [40]. In images, features with higher backscatter appear brighter, while those with lower backscatter appear darker. Water bodies present very dark features that contrast sharply with their surroundings due to their minimal roughness and consequently low backscatter, as illustrated in the figure below (Fig. 6). Water bodies were extracted using Sentinel-1 GRD SAR Data in ArcGIS Pro; with pixel classification performed using the Deep Learning tool available in the Image Analyst toolbox within ArcGIS Pro.

**Fig.6.**
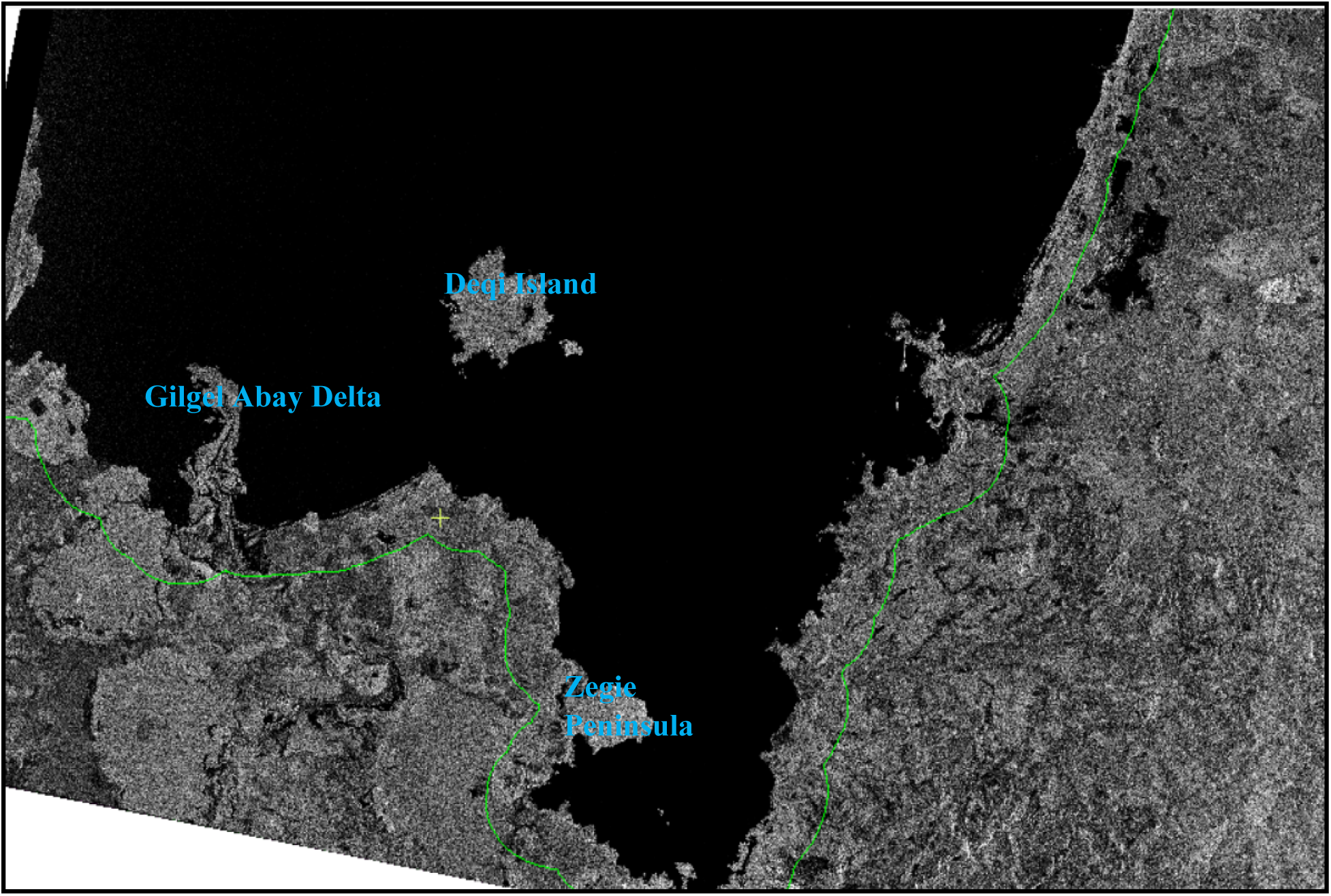
The difference in backscattering between the Gulf of Lake Tana region covered by water and other types of land cover: Water features appear as very dark features that contrast sharply with their surroundings land cover because they have very little roughness and, consequently, very low backscatter whereas the surrounding terrestrial parts including Zegie Peninsula, Deqi Island and Gilgel Abay Delta appear as white features because they have very high roughness and, consequently, very high backscatter.

#### Integrating Indicators for Shoreline Wetlands Mapping

In efforts to facilitate the mapping of shore area wetlands at Lake Tana, four key wetland indicators were selected: wetland hydrology, hydrophytic vegetation, hydric soil, and topographic position. Each of these indicators was converted into raster maps utilizing the same coordinate system (UTM zone 37N) and a uniform pixel size of 30m x 30m.

The Analytic Hierarchy Process (AHP) and the weighted overlay method were employed to combine these indicators for the mapping (equation 1). This study applied the AHP Python toolbox integrated within ArcGIS 10.7, performing pairwise comparisons as outlined by Saaty in the Analytical Hierarchy Method [41]. Initially, the AHP toolbox was added to ArcGIS 10.7, followed by the incorporation of AHP script tools. The AHP script was used to create an empty AHP matrix, which we then filled in (Table 1). The consistency index (CI) and consistency ratio (CR) were derived from the comparison matrix using the AHP script tool (Table 2).

**Table 1.**
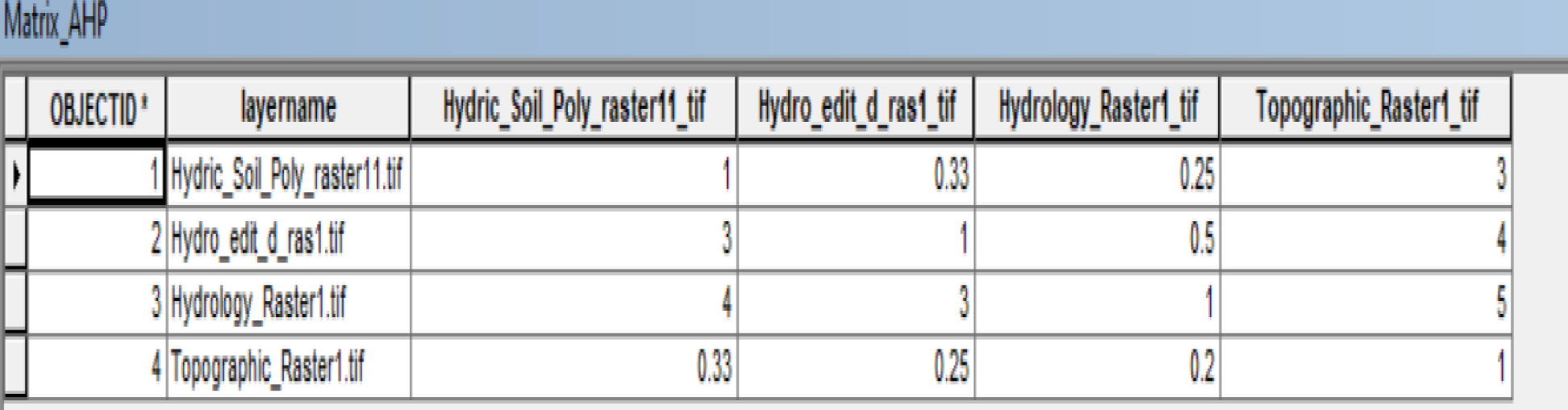
AHP matrix produced by AHP (step1) script and filled by the user

**Table 2.**
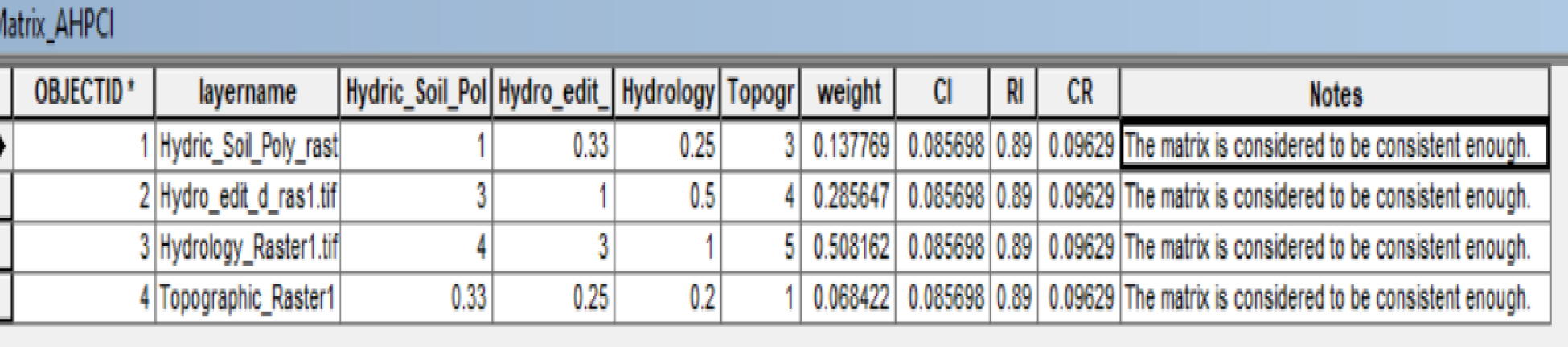
AHP matrix produced by AHP (step2) script that generate weight, consistency index (CI), the average consistency index (RI), consistency ratio (CR) and Notes

Subsequently, a comprehensive table was generated, detailing the weights assigned to each layer along with the consistency index (CI), consistency ratio (CR), and the average consistency index (RI). In this case, the CR (0.096) is below the threshold value of 1, indicating that the matrix is sufficiently consistent. The output raster could then be weighted according to importance and combined to produce a final output raster. The wetland indicators were assigned weights based on their contributions to wetland existence, with wetland hydrology recognized as the most critical variable among the three defining factors of wetlands: hydrology, hydric soil, and hydrophytic vegetation. This is because suitable hydrologic conditions primarily determine the other characteristics. As a result, the greatest weight was assigned to wetland hydrology, followed by hydrophytic vegetation.

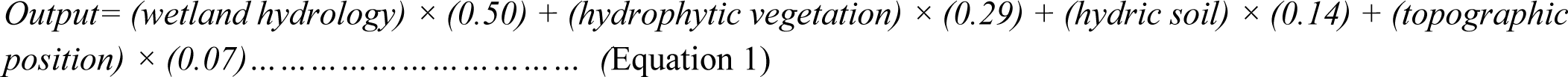

## Results

### Area Coverage of Shoreline Wetlands According to Individual Indicators

#### Topographic position

In Fig.7, the topographic position wetland indicator is illustrated, which was created by combining bathymetric data from depths of 0 to 2 meters with elevation maps ranging from 1786 to 1790 meters. This integration of data provides valuable insights into the ecological characteristics and spatial distribution of wetlands in this region. The topography position wetland indicator map covered about 55,363.53ha.

**Fig.7.**
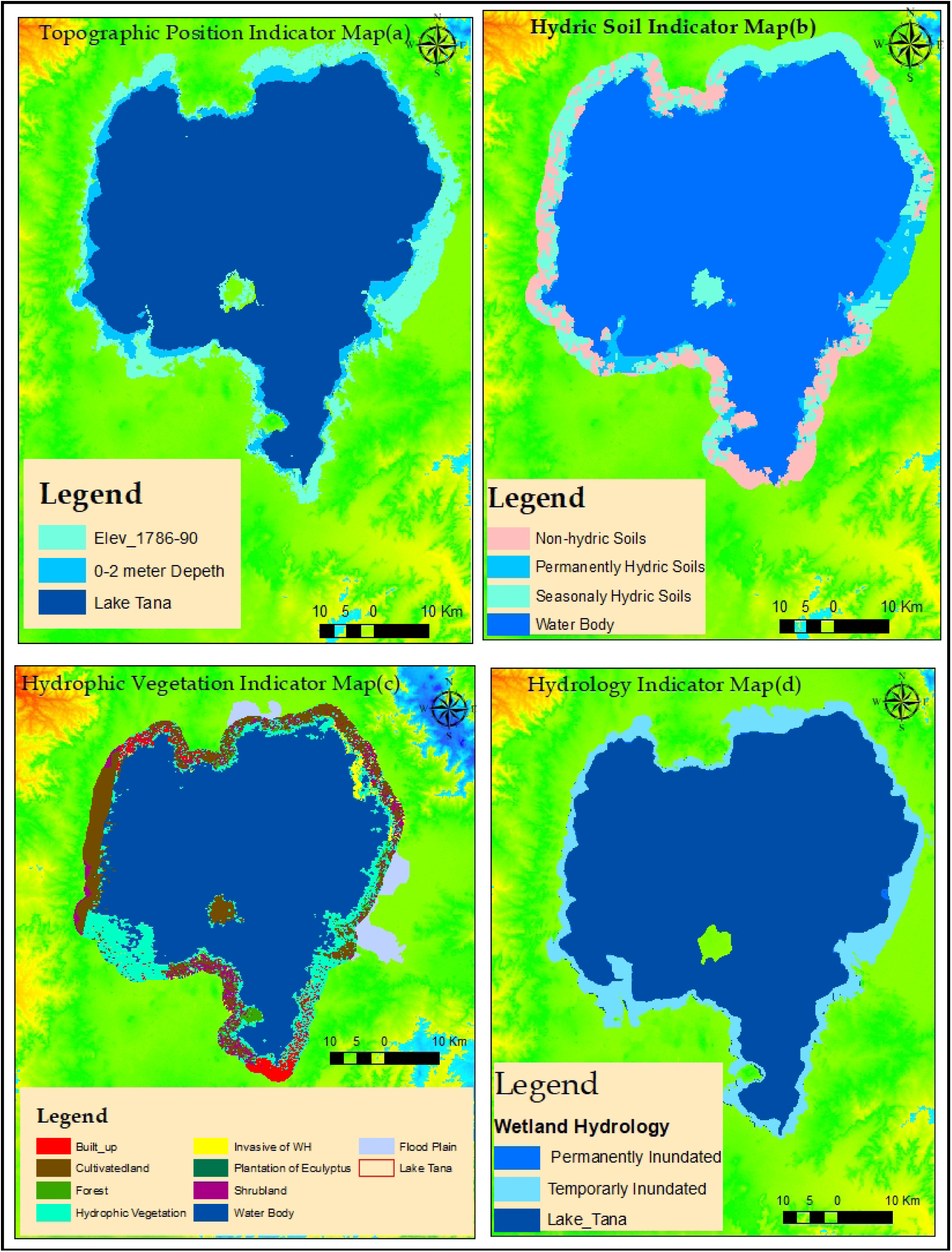
Shore area wetland indicator maps

#### Hydric Soil Indicators

According to the Bureau of Environmental Protection, Land Administration, and Use (2015), the soils within a 3 km buffer zone around Lake Tana consist of 12 major types. These include Vertisols (34,924.07 ha), Luvisols (8,693.89 ha), Nitisols (8,725.15 ha), Leptosols (7,708.54 ha), Alisols (6,105.11 ha), Cambisols (4,524.73 ha), Regosols (957.38 ha), Fluvisols (9,760.32 ha), Ferralsols (1,016.11 ha), Gleysols (10,466.38 ha), Acrisols (156.19 ha), and Lixisols (56.75 ha) (Fig.7b).

Based on recognized morphological characteristics, these soils were reclassified into hydric soils (including Vertisols, Fluvisols, Gleysols, and water) and non-hydric soils (including urban and built-up areas). Hydric soils were further categorized into water bodies, permanently hydric soils (Gleysols), and seasonally hydric soils (Vertisols and Fluvisols). Together, the permanently and seasonally hydric soils within the 3 km buffer zone of Lake Tana cover approximately 55,151 ha (Fig.7b).

Fluvisols are predominantly located in depressions along major streams and on lower, gently sloping plains. These soils are subject to frequent flooding and are continually replenished with new sediments during annual floods. They are primarily found in the Rib, Gilgel Abay, Gumara, and Shini rivers, as well as the Fogera plains, and are associated with floodplain and lacustrine temporary wetlands (Fig.7b). Gleysols, which have hydromorphic properties within 50 cm of the surface, form under prolonged wet conditions that create reducing environments. These conditions lead to the transformation of ferric compounds into ferrous compounds, resulting in hydric characteristics. Gleysols in Lake Tana are closely associated with lacustrine permanent wetlands and water depths of less than two meters. However, not all areas with hydric soils, including Vertisols, Fluvisols, and Gleysols, meet the criteria for classification as shore area wetlands. Some lack the hydrology typical of wetlands or do not support hydrophytic vegetation, which are essential characteristics of wetlands.

#### Hydrophytic Vegetation Indicators

Fig.7c illustrates eight major land use/cover classes around Lake Tana, derived from Sentinel-1A SAR input data and analyzed using the Object Analyst add-on within PCI Geomatica Banff software. The resulting map highlights the effectiveness of Object Analysts in identifying diverse land use/cover types within a 3 km buffer zone from the lake. These classifications were restructured into two primary categories: hydrophytic vegetation (including hydrophytic vegetation and invasive water hyacinth) and non-hydrophytic vegetation (comprising forests, built-up areas, cultivated land, shrub-land, Eucalyptus plantations, and water bodies).

The hydrophytic vegetation of shore area wetlands is widespread, except in the northern and northwestern parts of the lake (Fig.7c). Within the 3 km buffer, hydrophytic vegetation, including invasive water hyacinth, accounts for approximately 74,771.86 ha. Forests, covering 3,209.08 ha, are concentrated in areas such as Zegie, Kunzila, Woleta Petros, and Gorgora. Built-up areas, totaling 3,505.47 ha, are notable in Bahir Dar, Delgi, Kunzila, Sekelet, Sey-Deber, and Zegie.

The lake itself, along with cultivated land (18,939.38 ha), constitutes a significant portion of the area. Shrub-land, covering 8,434.68 ha, is prevalent in kebeles such as Abrehajerha, Agid Kirigna, Debranta, Dengel-Ber, Gobay Mariam, Kabe, Mangie, Merafit, Mitireha, Qorata, Tana Mistily, and Wagetera. Eucalyptus plantations (30.28 ha) encroach on wetland habitats in several kebeles, including Delgi, Estimut, Gorgora, Lijoma, Nabega, Zenzelima, Sekelet, Shum Abo, Tsizamba, Wagetera, and Weramit (Fig.7c).

The overall classification accuracy and Kappa statistic were measured at 78.68% and 0.70, respectively. Additionally, we computed the Producer Accuracy (PA), User Accuracy (UA), and Kappa Statistic (KS) for each class as shown in Table 3. The results revealed that water bodies achieved the highest accuracies (PA = 94.11%, UA = 99.12%, KS = 0.98), followed by cultivated land ((PA = 77.20%, UA = 81.04%, KS = 0.780). Hydrophytic vegetation showed moderate performance with PA at 69.69%, UA at 59.77%, and KS at 0.51. In contrast, shrub-land (PA = 45.00%, UA = 29.03%, KS = 0.27) and Eucalyptus plantations (PA = 0.00%, UA = 0.00%, KS = -0.02) exhibited the lowest PA, UA, and KS values (Table 3).

**Table 3.** Class Accuracy Statistics

| Class Name | Producer Accuracy (%) | User Accuracy (%) | Kappa Statistic |
| --- | --- | --- | --- |
| Hydrophytic vegetation | 69.69 | 59.77 | 0.51 |
| Forest | 58.62 | 45.13 | 0.44 |
| Built-up | 56.32 | 57.22 | 0.54 |
| Eucalyptus plantation | 0.00 | 0.00 | -0.02 |
| Waterbody | 94.11 | 99.12 | 0.98 |
| Cultivated land | 77.20 | 81.04 | 0.78 |
| Shrub-land | 45.00 | 29.03 | 0.27 |
| Water hyacinth | 54.55 | 58.21 | 0.56 |

**Table 4.** Area Coverage of Shoreline Wetlands According to Individual Indicators

| Wetland Indicator Type | Area(ha) |
| --- | --- |
| Topographic position | 55,364 |
| Hydric Soil | 55,151 |
| Hydrophytic Vegetation | 74,771 |
| Wetland Hydrology | 607,052 |

#### Wetland Hydrology

Using Sentinel-1 SAR data, two wetland hydrology categories were identified: (1) permanently inundated and (2) temporarily inundated areas. The hydrology wetland indicator map revealed that permanently inundated areas covered approximately 591,311.43 ha, while the inclusion of temporarily inundated areas increased the coverage to 607,052.48 ha (Fig.7d).

### Shore Area Wetlands Delineation Using Integrated Wetland Indicators

Shore area wetlands around Lake Tana extend approximately 3 km inland, covering 26,663.24 ha. The Fogera (19,576.64 ha) and Dembia (7,091.21 ha) floodplains constitute significant portions of these wetlands, located primarily in the eastern and northern parts of the lake (Fig. 8). These wetlands are found in topographic positions below 1790 meters and are characterized by either permanent or temporary inundation. They include areas with hydromorphic soils and hydrophytic vegetation, which are adapted to periodic flooding and specific wet conditions. Key wetland areas in the southern Gulf of Lake Tana include Aba-Garima, Selechen-Mariam, Avaji, Debo-Avanti, Giorgis-Alema, Shum-Abo, Gudguwad, and Gadero. Wetlands in the southwestern region encompass Dangel-Ber, Debranta, Estimut, the mouths of the Gilgel Abay and Infranz rivers, Kunzila, Lijoma-01, Lijoma-02, Sekelet, Sey-Deber, Wonjeta, and Yiganda-Zegie. In the southeast, wetlands such as Agid, Qirigna, Gileda, Bossit, the mouths of the Gumara and Rib rivers, Kabe, Merafit, Nabega, Qorata, Robit, Tana Mtsily, and Wagetera are prominent. Northern wetlands include Chemera, Chachna-Alwa, Mekonta-Ayibga, Fentaye-Narchacha, Delgi, Gorgora, the Dirma River Mouth, Tana-Woyna, Jerjer-Abanov, and Lemba-Arbaytu (Fig.8).

**Fig.8.**
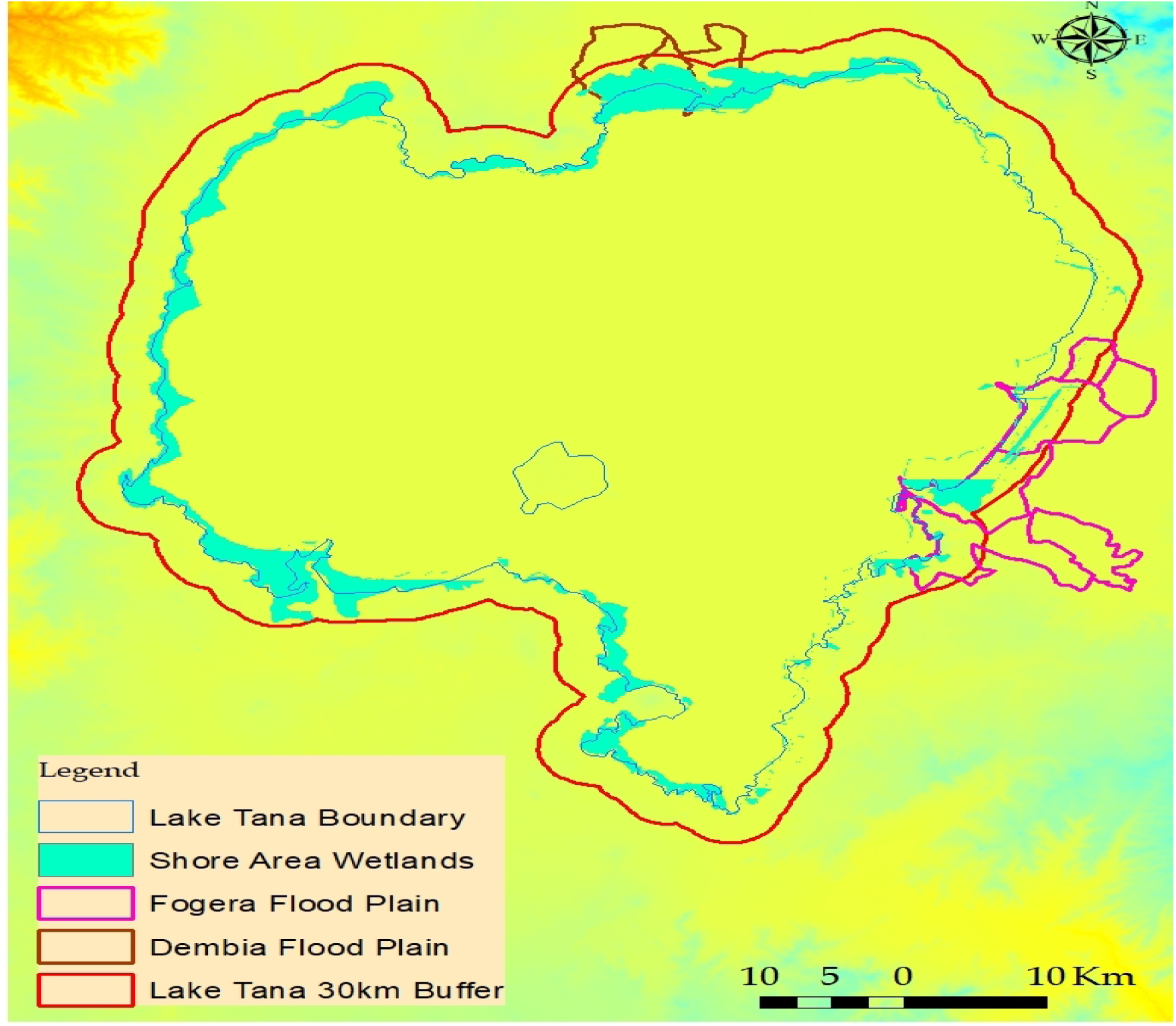
Shore area wetland in Lake Tana

Hydrophytic vegetation represents a critical structural element of the shore area wetland communities around Lake Tana. These wetlands can be classified into four groups based on the composition of hydrophytic vegetation (Fig.9):

1. **Southern Gulf of Lake Tana**: Dominated by reed macrophyte species.
2. **Northeastern Shore, Including the Fogera Floodplain**: Characterized by invasive water hyacinth macrophytes and a substrate of silt and sand sediments.
3. **Northern Dembia Floodplain**: Dominated by a combination of water hyacinth and *Echinochloa* spp., with silt as the primary sediment type.
4. **Northwestern Shore**: Predominantly composed of *Echinochloa* spp. and water hyacinth, with sand deposits as the dominant sediment type.

**Fig. 9.**
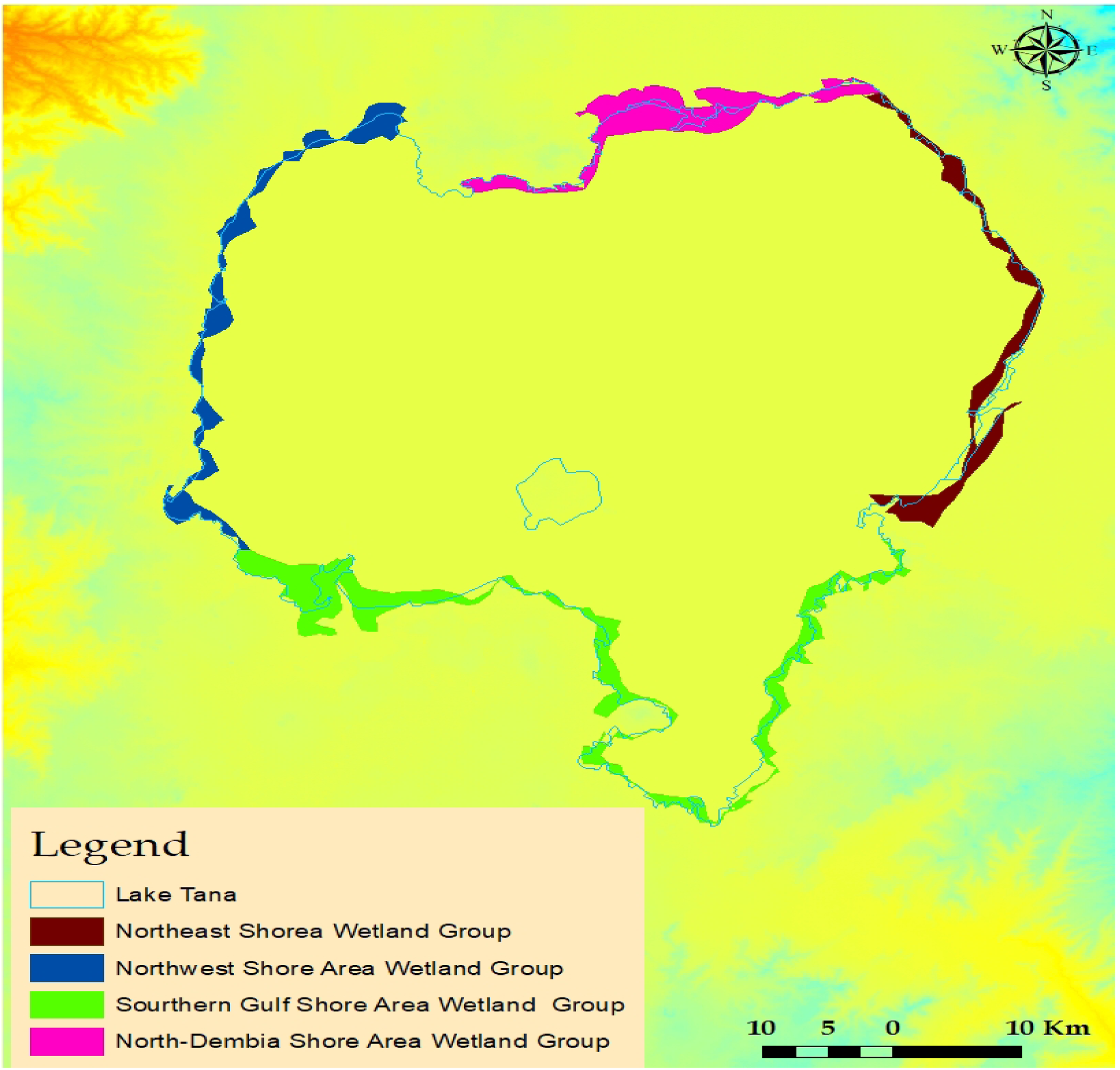
Shore area wetland groups in Lake Tana:

Southern Gulf of Lake Tana-dominated by reed (medium apple color);North Eastern, including Fogera flood plain-invaded by water hyacinth-silt and sand sediment type (dark brown color); North Dembia flood plain-dominated by water hyacinth and *Echinochloa spp.-silt* sediment type(pink color); and North Western-dominated *Echinochloa spp.* and water hyacinth-dominated by sand deposit (blue color).

## Discussion

Our study provides concrete evidence that Sentinel-1A SAR radar datasets significantly enhance shoreline wetland mapping, aligning with previous studies utilizing synthetic aperture radar (SAR) for wetland mapping [13–15,40,42–44]. In this research, we successfully mapped hydrophytic vegetation and wetland hydrology using publicly available Sentinel-1 SAR data processed with PCI Geomatica Banff software. Similarly, [40] employed multi-source Earth observation data, including RADARSAT-2 imagery processed through PCI Geomatica, for dynamic wetland mapping in the Great Lakes Basin. The use of SAR sensors, particularly Sentinel-1A, offers distinct advantages over optical sensors for shoreline wetland mapping. SAR can able to penetrate vegetation canopies and operate under all weather and illumination conditions, directly measuring the extent of flooded vegetation rather than inferring it from surface water extent [8]. Our findings align with prior studies indicating that multi-temporal SAR data enhances wetland classification [40,44,45].While SAR-based methods are effective, our results suggest that the integration of SAR with multi-source data, such as DEMs, can further improve classification accuracy, as highlighted by [43]. Despite these strengths, our classification of water hyacinth invasions yielded lower accuracy (PA = 54.55%, UA = 58.21%, KS = 0.56). Object-Based Image Analysis (OBIA) methods combined with high-resolution data, as used by [40] for mapping Great Lakes coastal wetlands, have demonstrated superior accuracy [46]. OBIA approaches could potentially be enhanced by incorporating SAR data into the classification process.

This study emphasizes the utility of Sentinel-1A SAR and PCI Geomatica Banff for shoreline wetland mapping, particularly under conditions where optical sensors are limited by cloud cover or lighting [15]. SAR’s ability to penetrate vegetation and soil provides valuable information unavailable to optical sensors [12,19]. However, sole reliance on SAR data may not always yield satisfactory accuracy. Integrating wetland indicators, such as hydric soils and topographic information derived from DEMs, can enhance mapping precision. Interestingly, our findings contrast with [42], who reported poor performance of Sentinel-1A SAR for herbaceous wetland mapping in the Biebrza floodplain (Poland), citing coarse resolution and limitations in detecting small wetland features. They achieved an overall accuracy (OAA) of 65% and a Kappa index (KIA) of 0.58 using multi-temporal SAR with VV/VH polarization. In comparison, our results demonstrated higher accuracy (OAA = 78.68%, KS = 0.70). Nonetheless, their findings highlighted the superior performance of TerraSAR-X/TanDEM-X images, especially in fully polarimetric mode.

Our classification results using Sentinel-1A SAR and PCI Geomatica showed good performance, with an overall accuracy of 78.68% and a Kappa coefficient of 0.70. These metrics are lower than those reported by [43] for wetland mapping in the Great Lakes (OA = 93.6%, K = 0.90), but higher than [40] 78% accuracy for targeted wetland classes. While our study accurately identified hydrophytic classes, including cultivated land and water bodies, non-hydrophytic classifications remained less precise. In contrast, [43] achieved higher accuracy for non-wetland classifications (OA = 96.62%, K = 0.95) compared to wetland classifications (OA = 87%, K = 0.91). Our classifier employed a support vector machine (SVM) method, which was also used by [42]. In contrast, [43] and [45] utilized the Random Forest (RF) classification algorithm, which has consistently been recognized as the most effective machine-learning approach for wetland mapping. RF classifiers have been particularly successful in categorizing wetlands into detailed classes, such as shallow water, marsh, swamp, fen, and bog, based on the Canadian Wetland Classification System [45].

The PCI Geomatica software used in this study supports various SAR sensors, including Sentinel-1, Radarsat-2, COSMO-SkyMed, TerraSAR-X, and others, offering extensive functionality for SAR data processing [40]. Geomatica’s Object Analyst, an advanced module, provides an integrated workflow for segmenting imagery, extracting features, building training sites, classifying data, refining shapes, and conducting accuracy assessments [35,47,48]. However, its accessibility is constrained by licensing and copyright restrictions, limiting its application for open-source data analysis. In comparison, platforms like Google Earth Engine (GEE) offer free access to satellite datasets and enable scalable wetland mapping without the need for extensive local storage [43,45,49].Launched in 2014, the Sentinel-1 satellite is the most recent SAR mission operating at the C-band and it’s freely available, which is another reason to explore the potential of this data for wetland InSAR applications [39].

The proposed method of integrating Sentinel-1A SAR with multi-source, multi-temporal data shows promise for generating annual and seasonal wetland maps in the Lake Tana Biosphere Reserve. Combining SAR data with spatial wetland indicators, such as hydric soils and topographic position improves classification accuracy and enables more precise monitoring of shoreline wetland dynamics.

## Conclusions

This study demonstrated that four wetland indicators (topographic position, hydric soil, hydrophytic vegetation, and hydrology) effectively map the spatial extent of shoreline wetlands in Lake Tana, northwestern Ethiopia. The success of this approach is largely attributed to the “double bounce” phenomenon, where radar pulses are backscattered twice, either from the water’s surface to the vegetation or vice versa, enhancing the detection of wetland features. From an overall classification accuracy, we could conclude that hydrophytic classes including water bodies and cultivated-land were identified more accurately than non-hydrophytic classes. This study concluded that SAR data alone could not produce satisfactory wetland classification accuracy. We minimize such limitations by integrating spatial geo-information indicators, such as topographic position and hydric soils derived from ArcGIS, with data captured by Sentinel-1A SAR, which focuses on hydrophytic vegetation and wetland hydrology, processed through PCI Geomatica Banff software. This combined approach ensures a more accurate and comprehensive representation of shoreline wetlands.

The results of this study provide a valuable baseline for future research and offer practical guidance for the sustainable management and conservation of shoreline wetland resources in the Lake Tana Biosphere Reserve. For managing and restoring shore area wetlands and making policy, the following measures are suggested: Shore area wetlands need to be delineated, and their buffer zone needs to be established using as a reference the results of this study.

### CRediT authorship contribution statement

Yirga K. Wondim: Conceptualization, Methodology, Data curation, Formal analysis, Software, Visualization and Writing – original draft**. Ayalew W. Melese:** Methodology, Funding acquisition, Project administration. Resources, Validation, Supervision and Writing – review & editing. **Workiyie W. Assefa:** Data curation, Visualization. Writing – review & editing, Supervision.

## Ethics declarations

### Ethical approval

All authors have read, understood, and have complied as applicable with the statement on “Ethical responsibilities of Authors” as found in the Instructions for Authors.

## Acknowledgment

We extend our sincere thanks to European Space Agency for providing access to Sentinel-1 Data product that were instrumental in advancing this research. The authors wish to thank the anonymous reviewers for their valuable comments and recommendations on the original version of this study.

## References

1. Chapman LJ, Baliirwa J, Bugwnyi FWB, Chapman C, Crisman TL. WETLANDS OF EAST AFRICA:BIODIVERSITY,EXPLOITATION,AND POLICY PERSPECTIVES. Biodivers Wetl,. 2001;2:101–131.

2. Leira M, Cantonati ÆM. Effects of water-level fluctuations on lakes : an annotated bibliography. 2008;171–84.

3. Vijverberg J, Sibbing FA, Dejen E. Lake Tana: Source of the Blue Nile. In: H.J. Dumont (ed.), editor. Springer Science + Business Media B.V. 2009; 2009. p. 163–92.

4. Krylov A V, Zelalem W, Prokin AA. Qualitative Composition and Quantitative Characteristics of Zooplankton in the Littoral Zone of Lake Tana (Ethiopia) at the End of the Dry Season. Inl Water Biol. 2020;13(2):206–13.

5. Wondie A. Ecological conditions and ecosystem services of wetlands in the Lake Tana Area, Ethiopia. Integr Med Res [Internet]. 2018; Available from: 10.1016/j.ecohyd.2018.02.002

6. Kassa Y, Mengistu S., Wondie A. TD. Distribution of macrophytes in relation to physico-chemical characters in the south western littoral zone of Lake Tana, Ethiopia - ScienceDirect. Aquat Bot [Internet]. 2021 [cited 2021 Nov 19];170(2021):1–9. Available from: https://www.sciencedirect.com/science/article/abs/pii/S0304377020301613

7. Adam E, Mutanga O, Rugege D. Multispectral and hyperspectral remote sensing for identification and mapping of wetland vegetation: A review. Wetl Ecol Manag. 2010;18(3):281–96.

8. Wu Q. GIS and Remote Sensing Applications in Wetland Mapping and Monitoring. 2018;(September 2017):140–57.

9. Rebelo L, Finlayson CM, Nagabhatla N. Remote sensing and GIS for wetland inventory, mapping and change analysis. J Environ Manage [Internet]. 2009;90(7):2144–53. Available from: 10.1016/j.jenvman.2007.06.027

10. Mwita E, Menz G, Misana S, Nienkemper P. Detection of Small Wetlands with Multi Sensor Data in East Africa. Adv Remote Sens. 2012;01(03):64–73.

11. Dubeau P, King DJ, Rebelo DGU and LM. Mapping the Dabus Wetlands, Ethiopia, Using Random Forest Classification of Landsat, PALSAR and Topographic Data. MDPI Remote Sens. 2017;9(1056):1–23.

12. Tiner RW, Lang MW, Klemas V V. Remote Sensing of Wetlands: Applications and Advances. Taylor & Francis Group; 2015. 566 p.

13. Kim JW. Applications of Synthetic Aperture Radar (SAR)/ SAR Interferometry (InSAR) for Monitoring of Wetland Water Level and Land Subsidence by. 2013;(503).

14. Wdowinski S, Hong S hoon. Wetland InSAR: a Review of the Technique and Applications. 2015;(March).

15. Lu Z, Member S, Kwoun O ig. Radarsat-1 and ERS InSAR Analysis Over Southeastern Coastal Louisiana : Implications for Mapping Water-Level Changes Beneath Swamp Forests. 2008;46(8):2167–84.

16. Hong S hoon, Wdowinski S, Kim S wan, Won J sun. Remote Sensing of Environment Multi-temporal monitoring of wetland water levels in the Florida Everglades using interferometric synthetic aperture radar (InSAR). Remote Sens Environ [Internet]. 2010;114(11):2436–47. Available from: 10.1016/j.rse.2010.05.019

17. Richards JA. An explanation of enhanced radar backscattering from flooded forests. 1987;8(7):1093–100.

18. Mahdavi S, Salehi B, Amani M, Granger JE, Brisco B, Huang W, et al. Object-based Classification of Wetlands in Newfoundland and Labrador Using Multi-Temporal PolSAR Data. 2017 8992(June).

19. Salehi B, Mahdianpari M, Amani M, Manesh FM, Granger J, And SM, et al. Wetlands Management - Assessing Risk and Sustainable Solutions. In: Wetlands Management - Assessing Risk and Sustainable Solutions. 2018. p. 110.

20. Mahdianpari M, Salehi B. A new speckle reduction algorithm of polsar images based on a combined Gaussian random field model and wavelet edge detection approach A NEW SPECKLE REDUCTION ALGORITHM OF POLSAR IMAGES BASED ON A COMBINED GAUSSIAN RANDOM FIELD MODEL AND WAVELET EDGE DETE. 2017;(January 2018).

21. Guo M, Li J, Sheng C, Xu J, Wu L. A review of wetland remote sensing. Sensors (Switzerland). 2017;17(4).

22. Gondwe BRN, Hong S hoon. Hydrologic Dynamics of the Ground-Water-Dependent Sian Ka ’ an Wetlands, Hydrologic Dynamics of the Ground-Water-Dependent Sian Ka ’ an Wetlands, Mexico, Derived from InSAR and SAR Data. 2014;(February 2010).

23. Dagne SS, Hirpha HH, Tekoye AT, Dessie YB. Fusion of sentinel – 1 SAR and sentinel – 2 MSI data for accurate Urban land use – land cover classification in Gondar City, Ethiopia. Environ Syst Res [Internet]. 2023;5. Available from: 10.1186/s40068-023-00324-5

24. Wondim YK, Melese AW. Evaluation of the evapotranspiration rate of lacustrine wetland macrophytes in Lake Tana, Ethiopia. Ecohydrol Hydrobiol [Internet]. 2023;23(4):623–34. Available from: 10.1016/j.ecohyd.2023.05.003

25. Alemayehu T, Mccartney M, Kebede S. The water resource implications of planned development in the Lake Tana catchment, Ethiopia. Ecohydrol Hydrobiol. 2010 Jan 1;10(2–4):211–21.

26. Kebedew MG, Tilahun SA, Zimale FA, Belete MA, Wosenie MD, Steenhuis TS. Relating Lake Circulation Patterns to Sediment, Nutrient, and Water Hyacinth Distribution in a Shallow Tropical Highland Lake. Hydrology. 2023;10(9).

27. Smith RD, Ammann A, Bartoldus C, Brinson MM. US Army Corps of Engineers Wetlands Research Program Technical Report WRP-DE-9. 1995.

28. Tiner RW. Defining Hydrophytes for Wetland Identification and Delineation ERDC / CRREL CR-12-1 Defining Hydrophytes for Wetland Identification and Delineation Cold Regions Research and Engineering Laboratory. 2015;(January 2012).

29. Huertos ML, Smith D. Wetland Bathymetry and Mapping. In 2013. p. 38.

30. Kebedew MG, Tilahun SA, Belete MA, Zimale FA, Steenhuis TS. Sediment deposition (1940–2017) in a historically pristine lake in a rapidly developing tropical highland region in Ethiopia. Earth Surf Process Landforms. 2021;46(8):1521–35.

31. Tiner RW. WETLAND INDICATORS : A GUIDE TO WETLAND IDENTIFICATION, DELINEATION, CLASSIFICATION, AND MAPPING. New York: CRC Press LLC CRC; 1999. 399 p.

32. Ali A, Erkossa T, Gudeta K, Abera W, Mesfin E, Mekete T, et al. Reference soil groups map of Ethiopia based on legacy data and machine learning-technique: EthioSoilGrids 1.0. Soil. 2024;10(1):189–209.

33. Bureau of Environmental Protection LA and U. Amhara National Regional State. Vol. I. Bahir Dar; 2015.

34. Tiner RW. Ecology of Wetlands: Classification Systemss. In: Encyclopedia of Inland Waters [Internet]. 2009. p. 516–25. Available from: https://linkinghub.elsevier.com/retrieve/pii/B9780123706263000570

35. Shah Hosseini R, Entezari I, Homayouni S, Motagh M, Mansouri B. Classification of polarimetric SAR images using Support Vector Machines. Can J Remote Sens. 2011;37(2):220–33.

36. Sukawattanavijit C, Jie C. GA-SVM Algorithm for Improving Land-Cover Classification Using SAR and Optical Remote Sensing Data. IEEE Geosci Remote Sens Lett. 2017;14(3):284–8.

37. Mantero P, Moser G, Serpico SB. Partially supervised classification of remote sensing images using SVM-based probability density estimation. 2003 IEEE Work Adv Tech Anal Remote Sensed Data. 2004;43(3):327–36.

38. Story M, Congalton RG. Remote Sensing Brief Accuracy Assessment: A User’s Perspective. Photogramm Eng Remote Sensing [Internet]. 1986;52(3):397–9. Available from: https://www.asprs.org/wp-content/uploads/pers/1986journal/mar/1986_mar_397-399.pdf

39. Mohammadimanesh F, Mahdianpari M, Salehi B, Brisco B. Wetland Water Level Monitoring Using Interferometric Synthetic Aperture Wetland Water Level Monitoring Using Interferometric Synthetic Aperture Radar (InSAR): A Review. Can J Remote Sens [Internet]. 2018;0(0):1–16. Available from: 10.1080/07038992.2018.1477680

40. Battaglia MJ, Banks S, Behnamian A, Bourgeau-Chavez L, Brisco B, Corcoran J, et al. Multi-source EO for dynamic wetland mapping and monitoring in the great lakes basin. Remote Sens. 2021;13(4):1–38.

41. Saaty TL. How to make a decision : The Analytic Hierarchy Process. 1990;48.

42. Mleczko M, Mroz M. Wetland Mapping Using SAR Data from the Sentinel-1A and TanDEM-X Missions : A Comparative Study in the Biebrza Floodplain (Poland). Remote Sens. 2018;10(78):1–19.

43. Mohseni F, Amani M, Mohammadpour P, Kakooei M, Jin S, Moghimi A. Wetland Mapping in Great Lakes Using Sentinel-1/2 Time-Series Imagery and DEM Data in Google Earth Engine. Remote Sens. 2023;15(14).

44. Banks S, Millard K, Behnamian A, White L, Ullmann T, Charbonneau F, et al. Contributions of Actual and Simulated Satellite SAR Data for Substrate Type Differentiation and Shoreline Mapping in the Canadian Arctic. Remote Sens. 2017;9(1206):27.

45. Amani M, Mahdavi S, Afshar M, Brisco B, Huang W, Mohammad S, et al. Canadian Wetland Inventory using Google Earth Engine : The First Map and Preliminary Results. 2019;1–20.

46. Dronova I. Object-Based Image Analysis in Wetland Research: A Review. 2015;6380–413.

47. Blaschke T, Hay GJ, Kelly M, Lang S, Hofmann P, Addink E, et al. Geographic Object-Based Image Analysis - Towards a new paradigm. ISPRS J Photogramm Remote Sens [Internet]. 2014;87:180–91. Available from: 10.1016/j.isprsjprs.2013.09.014

48. Hossain MD, Chen D. Segmentation for Object-Based Image Analysis (OBIA): A review of algorithms and challenges from remote sensing perspective. ISPRS J Photogramm Remote Sens [Internet]. 2019;150(February):115–34. Available from: 10.1016/j.isprsjprs.2019.02.009

49. Gulácsi A, Ferenc K. Sentinel-1-Imagery-Based High-Resolution Water Cover Detection on Wetlands, Aided by Google Earth Engine. 2020;1–20.

